# PKCδ splicing variant (PKC*δ*III) induces neurite outgrowth in PC12 cells in the absence of NGF

**DOI:** 10.1101/2023.08.23.554440

**Authors:** Hiroyuki Yamasaki, Tomoko Nakai, Kanae Kitatani, Chikara Kato, Susumu Takekoshi

**Affiliations:** Department of Cell Biology, Division of Host Defense Mechanism, Tokai University School of Medicine, Isehara 259-1193, Japan; Department of Molecular Life Science, Tokai University School of Medicine, Isehara 259-1193, Japan; Medical Science College Office, Tokai University School of Medicine, Isehara 259-1193, Japan; College of Agriculture, Academic Institute, Shizuoka University, Shizuoka 422-8529, Japan

**Author notes:** Corresponding author: Susumu Takekoshi, Department of Cell Biology, Division of Host Defense Mechanism, Tokai University School of Medicine. Author contributions: HY, KK, CK, and ST designed the study; HY and TN performed the experiments; HY, TN, and ST analyzed data; HY and TN wrote the paper; HY, TN, KK, CK, and ST edited the paper.

**Keywords:** PKCδ, neurite outgrowth, splicing variant, PKC*δ*III

## Abstract

Protein kinase C (PKC)δ is a serine/threonine kinase involved in many cellular processes in response to diverse stimuli, including cell proliferation, differentiation, apoptosis, tumor inhibition, and cell migration. PKCδ consists of the N terminal regulatory domain and the C terminal catalytic domain, which are linked with a hinge region. An alternative splicing variant of rat PKCδ (also known as PKCδIII) has been reported, and this PKCδ splicing variant (SV) has only the regulatory domain without the catalytic domain. However, its function remains unclear. In this study, we found that PKCδ SV induced neurite outgrowth in the absence of nerve growth factor (NGF) in PC12 cells. Immunoblot analysis revealed that ERK was phosphorylated by NGF treatment, but not by PKCδ SV induction, indicating that PKCδ SV-mediated neurite outgrowth was different from NGF-mediated one. In addition, we showed that PKCδ full length and SV increased their mRNA expression after NGF treatment and that PKCδ SV was more susceptible to its proteasomal degradation. Our findings provided insights into the role of PKC*δ* SV in neurite outgrowth.

## Introduction

Neurons differentiate from spherical shaped cells to highly polarized cells with axons and dendrites. Neurites, the precursors of axons and dendrites, emerge from the cell body and develop during neuronal migration to establish the proper neuronal network [1–3]. Dysregulation of neurite outgrowth and neuronal migration have harmful effects on the synaptic connections, which result in neurodevelopmental disorders such as autism, schizophrenia, lissencephaly, and microcephaly [3,4].

Protein kinase C (PKC) is a phospholipid-dependent family of serine/threonine protein kinases. There are eleven PKC isoforms that are subdivided into three groups: conventional PKCs (α, βI, βII and γ), novel PKCs (δ, ε, η and θ), and atypical PKCs (ζ and λ/ι). Conventional PKCs are activated by diacylglycerol (DAG) in a calcium-dependent manner. In contrast, novel PKCs are activated by DAG in a calcium-independent manner. Atypical PKCs activation does not require DAG nor calcium. PKCδ belongs to the novel PKC family, and is involved in many cellular processes in response to diverse stimuli, including cell proliferation, differentiation, apoptosis, tumor inhibition, and cell migration [5].

PKCδ ablation in mice affected arteriosclerosis [6], B cell tolerance [7,8], insulin signaling [9], and fertility [10]. A recent report demonstrated that PKCδ KO mice lacking different exons result in the different outcomes [11]. While PKCδ KO mice lacking the second exon develop normally and are fertile [6], PKCδ KO mice lacking exon 7 show short life span and are not born in accordance with Mendelian rules [11]. These reports highlight the importance of remaining PKCδ transcript variants in PKCδ KO mice.

There are 9 splicing variants in humans, mice and rats. In this study, we focused on PKCδ splicing variant (SV) found in rats, whose function remains uncharacterized [12]. Therefore, we sought to elucidate the function of this PKCδSV in rat adrenal pheochromocytoma PC12 cells. PC12 cells are widely used as a model for neurite outgrowth. Moreover, the regulatory domain (RD) of PKC*δ*, which covers almost all sequences of PKC*δ* SV, induces neurite outgrowth in rat neural cells [13]. Because only PKC*δ* RD cannot be expressed *in vivo*, we explored the relationship between PKCδ SV and neurite outgrowth.

In this study, we showed that PKCδ SV induced neurite outgrowth in the absence of nerve growth factor (NGF). Our data also demonstrated that PKCδ SV increased after NGF treatment in mRNA level and that PKCδ SV was more susceptible to its proteasomal degradation than other PKCδ proteins were. Our findings provided insights into the role of PKC*δ* SV in neurite outgrowth.

## Materials and methods

### Cell culture

PC12 cell line was purchased from the American Type Culture Collection (ATCC; Manassas, VA, USA). GP2-293 cell line was purchased from (Takara Bio, Kusatsu, Shiga, Japan). PC12 cells were cultured in RPMI-1640 with 2mM L-glutamine (FUJIFILM Wako Pure Chemical, Osaka, Japan) supplemented with 10% heat-inactivated horse serum (HS; Cambrex, East Rutherford, NJ, USA) and heat-inactivated 5% fetal bovine serum (FBS; Sigma-Aldrich, St. Loius, MO, USA) at 37°C in a humidified 5% CO_2_ incubator. GP2-293 cells were cultured in DMEM (FUJIFILM Wako Pure Chemical) supplemented with 10% tetracycline-free FBS (Takara Bio).

### Plasmids, reagents and retrovirus production

EGFP-tagged rat PKCδ cDNA were cloned into EcoRI and BamHI digested pRetroX-TetOne-Puro (#634307, Takara Bio) using In-Fusion cloning system (#639648, Takara Bio). All constructs were verified by direct sequencing. To produce retroviruses, pRetroX-TetOne-Puro-PKCδ constructs were cotransfected into GP2-293 cells with pVSV-G using Xfect Transfection Reagent. Supernatants were collected at 72h after transfection. After virus transduction, cells were cultured for at least 48h before selection was performed using 2μg/ml puromycin.

Other reagents included doxycycline hyclate (D9891, Sigma-Aldrich), MG-132 (S2619, Selleck Chemicals, Houston, TX, USA), DMSO (D8418, Sigma-Aldrich), β-nerve growth factor (AF-450-01-100μg, PeproTech, Cranbury, NJ, USA), TRITC conjugated phalloidin (H-1600, Vector laboratories, Newark, CA, USA), and HardSet Antifade Mounting Medium with DAPI (H-1500, Vector laboratories).

### Neurite extension assay

To evaluate PKCδ RD- and SV-mediated neurite outgrowth, cells were cultured in collagen coated chamber slides at a density of approximately 6x10^3^ cells/cm^2^. The following day, the medium was replaced with RPMI-1640 supplemented with 1% HS and 1μg/ml doxycycline hyclate (Dox, Sigma-Aldrich). Another 48h later, cells were fixed with 10% formaldehyde and permeabilized with 0.1% Triton X-100. After blocking with 3% BSA, cells were stained with TRITC conjugated phalloidin (H-1600, Vector laboratories) and mounted with HardSet Antifade Mounting Medium with DAPI (H-1500, Vector laboratories). Images were captured using Axio Imager M2 (Zeiss, Baden-Wurttemberg, Germany).

Neurite extension was analyzed using Image J (version 1.53s, downloaded on 19 May 2022, National Institutes of Health, Bethesda, MD, USA) software. Neurites were measured if the projection was 2μm or more in length, otherwise scored as zero. When a single cell possessed two or more neurites (>2μm, each), total length of neurites per cell was scored. At least 200 EGFP-positive cells per each condition were observed and the total neurites length were measured. Cells with neurites of 5μm or more were determined as neurite-positive cells.

To evaluate NGF mediated neurite outgrowth, PC12 cells were seeded at a density of 3.6x104/cm2 on collagen-coated 6well plates. The following day, the medium was replaced with RPMI-1640 supplemented with 1% HS and 100ng/ml *β*-NGF. Media containing NGF was exchanged every second day. Cells were harvested at the indicated time points for further analyses.

### Immunoblotting

Cells were washed with cold 0.15M NaCl and precipitated with 10% trichloroacetic acid on ice for 30min. Precipitates were lysed in cell lysis buffer (20 mM Tris/HCl, pH7.5; 1xLithium Dodecyl Sulfate sample buffer (Thermo Fisher Scientific, Waltham, MA, USA); a protease inhibitor cocktail (cOmplete, Merck, St. Louis, MO, USA); a phosphatase inhibitor cocktail (PhosSTOP, Merck). The samples were boiled, resolved by SDS-PAGE, and transferred to nitrocellulose membranes. The primary antibodies used for immunoblotting were as follows: anti-C-terminus of PKCδ (ab182126, 1:5000; Abcam, Cambridge, England), anti-N-terminus of PKC δ (#9616, 1:500; CST), anti-phospho-PKCδ/θ, Ser643/676 (#9376, 1:800; CST), anti-Ubiquitin (#58395, 1:1000; CST, Danvers, MA, USA), anti-GFP (PABG1, 1:1000; Proteintech, Rosemont, IL, USA), and anti-GAPDH (G9545, 1:5000; Sigma-Aldrich).

### qRT-PCR

Total RNA was isolated using TRIzol reagent (#15596026, Thermo Fisher Scientific). For each sample, 2 μg of total RNA was reverse transcribed into cDNA using a High-Capacity RNA-to-cDNA kit (#4387406, Applied Biosystems, Waltham, MA, USA) and sequences were amplified using Taq DNA polymerase that had been supplied with a PCR master mix (#4324018, Applied Biosystems). Relative quantification of the target mRNA was performed using the comparative CT method with the endogenous control glyceraldehyde-3-phosphate dehydrogenase (GAPDH) provided as predeveloped TaqMan Assay Reagents by Applied Biosystems. Assay IDs of Taqman probes for rat *Prkcd, Prkcd3*, and *Gapdh* were Rn00440891_m1, AJHSOQU (Custom Plus Taqman RNA Assays), and Rn01775763_g1, respectively. The PCR amplification and analysis were performed on a QuantStudio 3 real-time PCR instrument (Applied Biosystems).

### Statistical analysis

Data are presented as mean±standard deviation (SD). Statistical significance of differences was analyzed using one-way ANOVA. Dunnett’s test or Bonferroni’s test were performed as post-hoc test.

## Results

PKCδ consists of the N terminal regulatory domain and the C terminal catalytic domain, which are linked with a hinge region (Fig. 1A). PKCδ is proteolytically cleaved by caspase-3 at the hinge region. The cleavage of PKCδ allows catalytic domain to dissociate from the inhibitory regulatory domain and show its kinase activity. PKCδ splicing variant (SV) reported in rats lacks catalytic domain, and possesses only the inhibitory regulatory domain [12] (Fig. 1A). Given that the regulatory domain of PKCδ induces neurite outgrowth in rat neural cells [13], we hypothesized that the expression of PKCδ SV also induces neurite outgrowth. To test this possibility, we established doxycycline-inducible various PKCδ expressing PC12 cells (Fig. 1A). Although various PKCδ proteins were induced after doxycycline addition, the protein levels of truncated PKCδ proteins were lower than that of PKCδ full length (FL) (Fig. 1B-D).

**Figure 1.**
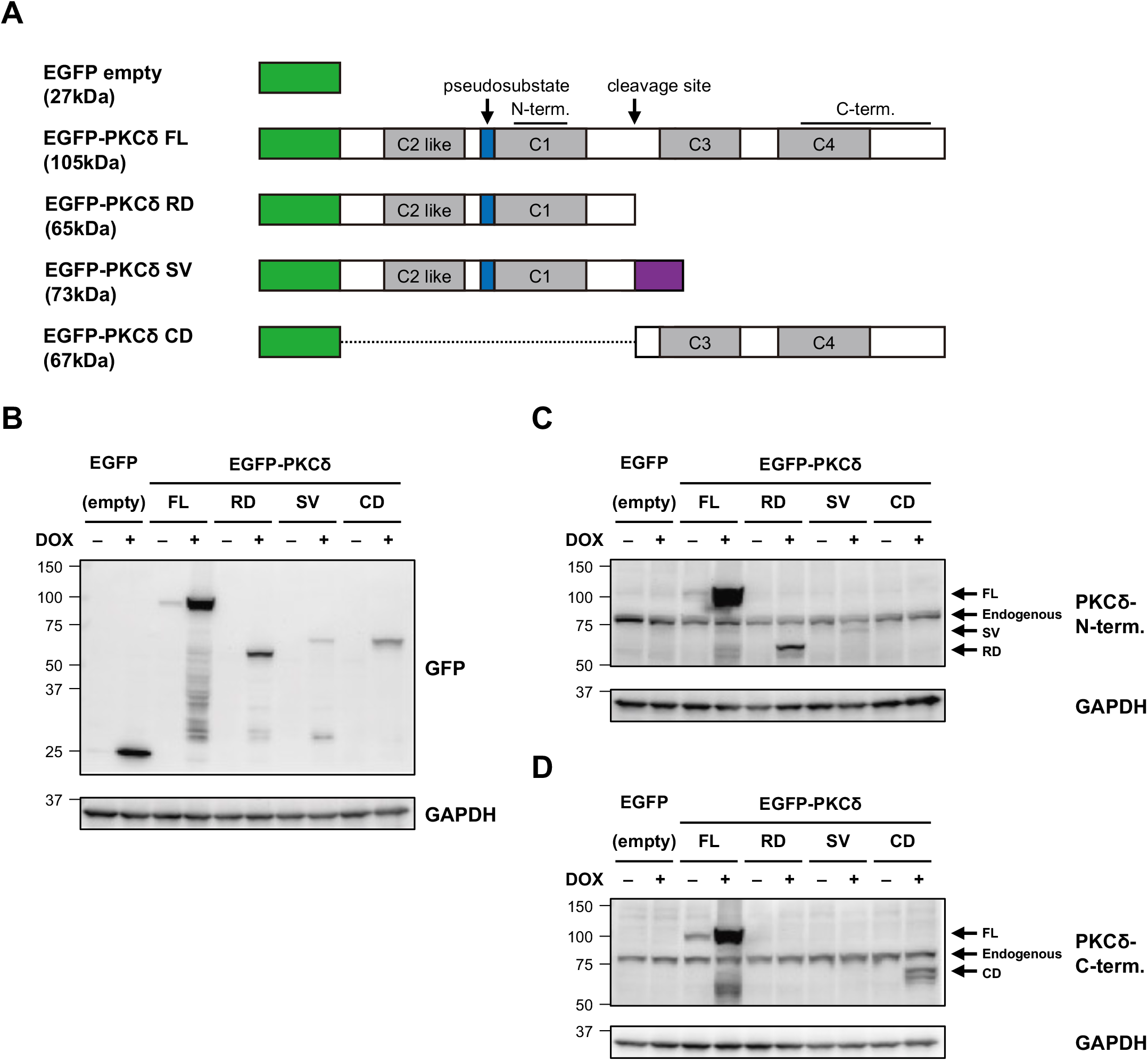
Establishment of PC12 cells containing various PKCδ constructs. (A) Schematics of PKCδ constructs used in this study. Black bars in PKCδ full length indicate the sequences that antibodies recognize. (B-D). Induction of PKCδ constructs. PC12 cells containing the indicated PKCδ constructs were cultured with or without 1ug/ml doxycycline for 24h. The cell lysates were subjected to immunoblotting analysis using the antibodies against GFP (B), PKCδ (N-term) (C), and PKCδ (C-term) (D). GAPDH was used as a loading control.

Consistent with the previous report [13], the expression of PKCδ regulatory domain (RD) induced neurite outgrowth in the absence of nerve growth factor (NGF) in PC12 cells (Fig. 2A, B and C). The expression of PKCδ SV also induced neurite outgrowth, while those of EGFP and the other PKCδ proteins did not (Fig. 2A, B, and C). These data provided the evidence that PKCδ SV, like PKCδ RD, induces neurite outgrowth in the absence of NGF. MAPK pathway plays a pivotal role in neurite outgrowth induced by NGF [14,15]. PKCδ is translocated to membrane after NGF treatment [16] and involved in NGF-mediated neurite outgrowth and ERK1/2 phosphorylation [17]. As expected, ERK1/2, a molecule of MAPK pathway, were phosphorylated after NGF treatment (Fig. 2D). On the other hand, PKCδ was phosphorylated even before NGF treatment and its phosphorylation was down-regulated by NGF treatment (Fig. 2D). This observation prompted us to test the possibility that exogenous PKCδ RD and SV may induce the phosphorylation of ERK1/2 and activate MAPK pathway with suppressing PKCδ phosphorylation. To this end, we examined the impact of exogenous various PKCδ proteins on endogenous PKCδ and ERK1/2 phosphorylation. Surprisingly, ERK1/2 was not phosphorylated after all PKCδ proteins induction (Fig. 2E). PKCδ phosphorylation was not influenced by all PKCδ proteins induction neither. These findings indicated that PKCδ RD and SV induce neurite outgrowth in an ERK-independent manner without influencing endogenous PKCδ. Furthermore, these findings also indicated that PKCδ RD- and SV-mediated neurite outgrowth was different from NGF-mediated neurite outgrowth.

**Figure 2.**
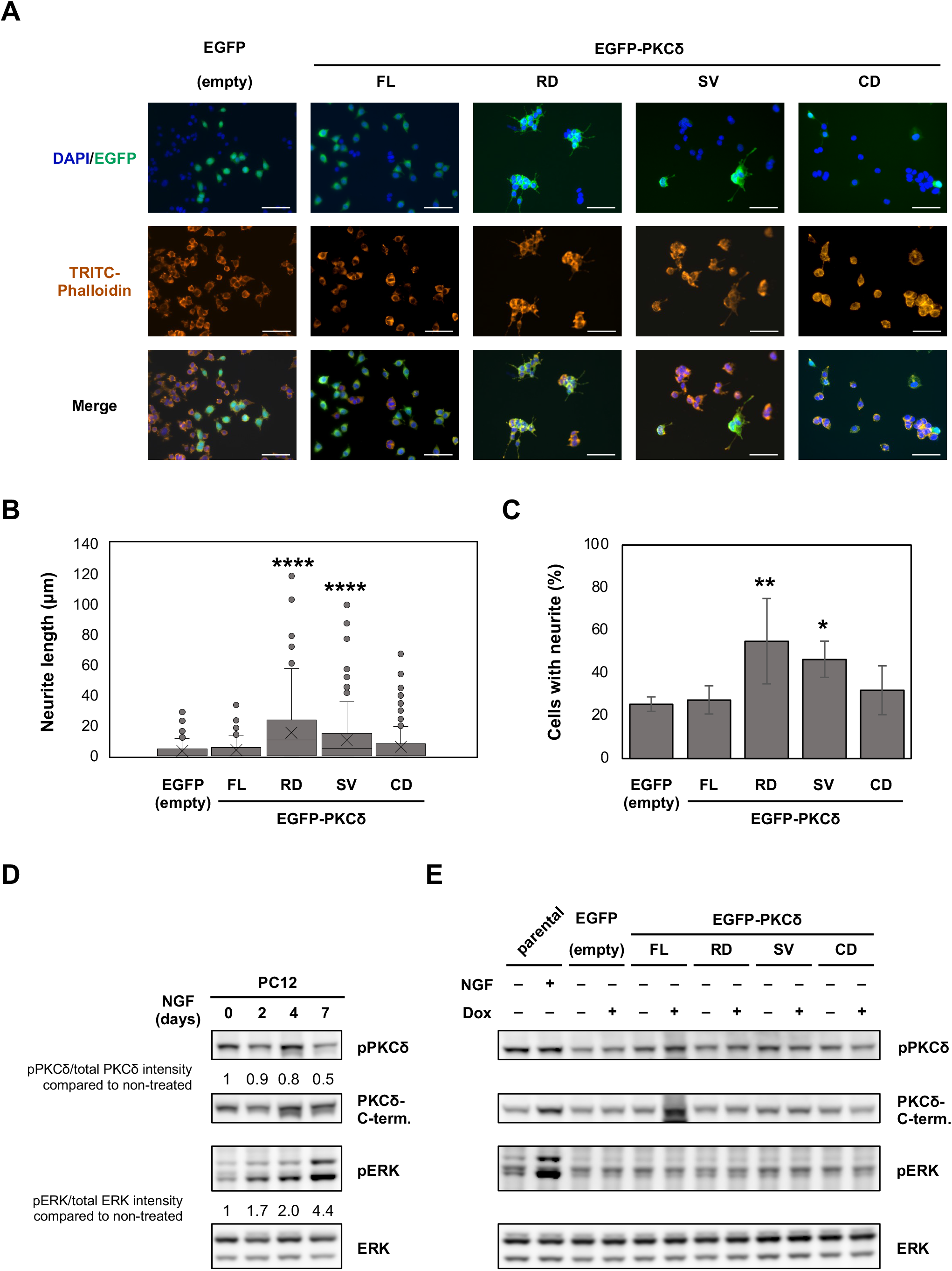
PKC*δ* SV induces neurite outgrowth in the absence of NGF in an ERK-independent manner. (A) Representative fluorescent images of PC12 cells containing the indicated EGFP-tagged PKCδ proteins. Cells were cultured with or without 1ug/ml doxycycline for 48h. After fixation, DNA was stained with DAPI and actin filament was stained with TRITC conjugated phalloidin. Scale bar represents 50μm. (B) Boxplots of total neurite length per a GFP-positive cell among PC12 cells containing the indicated PKCδ constructs. **** *p*<0.001 (one-way ANOVA [Dunnett’s test]). (C) Quantification of cells with >5um neurites in GFP-positive cells. Graph represent mean ± SD of *n*=5. * *p*<0.05; ** *p*<0.01 (one-way ANOVA [Dunnett’s test]). (D) PC12 cells were cultured with or without 100ng/ml NGF. After 2, 4, and 7 days, whole cell lysates were harvested. The cell lysates were subjected to immunoblotting analysis using the indicated antibodies. *n* = 3. (E) PC12 cells containing PKCδ constructs were cultured with 1ug/ml doxycycline. PC12 cells were also cultured with or without 100ng/ml NGF. 48h later, cells were harvested and the whole cell lysate were subjected to immunoblotting analysis using the indicated antibodies. *n* = 3.

To further assess the importance of PKCδ SV in neurite outgrowth, we monitored the expression of endogenous PKCδ FL and SV in both mRNA and protein levels during NGF-mediated neurite outgrowth (Fig. 3A). NGF treatment upregulated mRNA expression of both PKCδ FL and SV at day 4 (Fig. 3B). We sought to detect PKCδ FL and SV with two antibodies binding to the N terminus and the C terminus of PKCδ (Fig. 1A). Surprisingly, while PKCδ FL detected by immunoblotting using anti-PKCδ (C term) increased during NGF treatment (Fig. 3C, right panel), PKCδ FL detected by immunoblotting using anti-PKCδ (N term) did not correspond (Fig. 3C, left panel). This observation implies that the region recognized by anti-PKCδ (N term) might be modified by NGF treatment, resulting in a reduced affinity to anti-PKCδ (N term). In addition, NGF treatment induced PKCδ cleavage and the catalytic fragment was detected (Fig. 3C, right panel), indicating that NGF activates PKCδ signaling. In contrast, although PKCδ SV increased after the treatment with NGF in mRNA level, PKCδ SV was not detected by immunoblotting using anti-PKCδ (N term) (Fig. 3C, left panel). These results suggested that NGF treatment regulates PKCδ SV expression in mRNA level.

**Figure 3.**
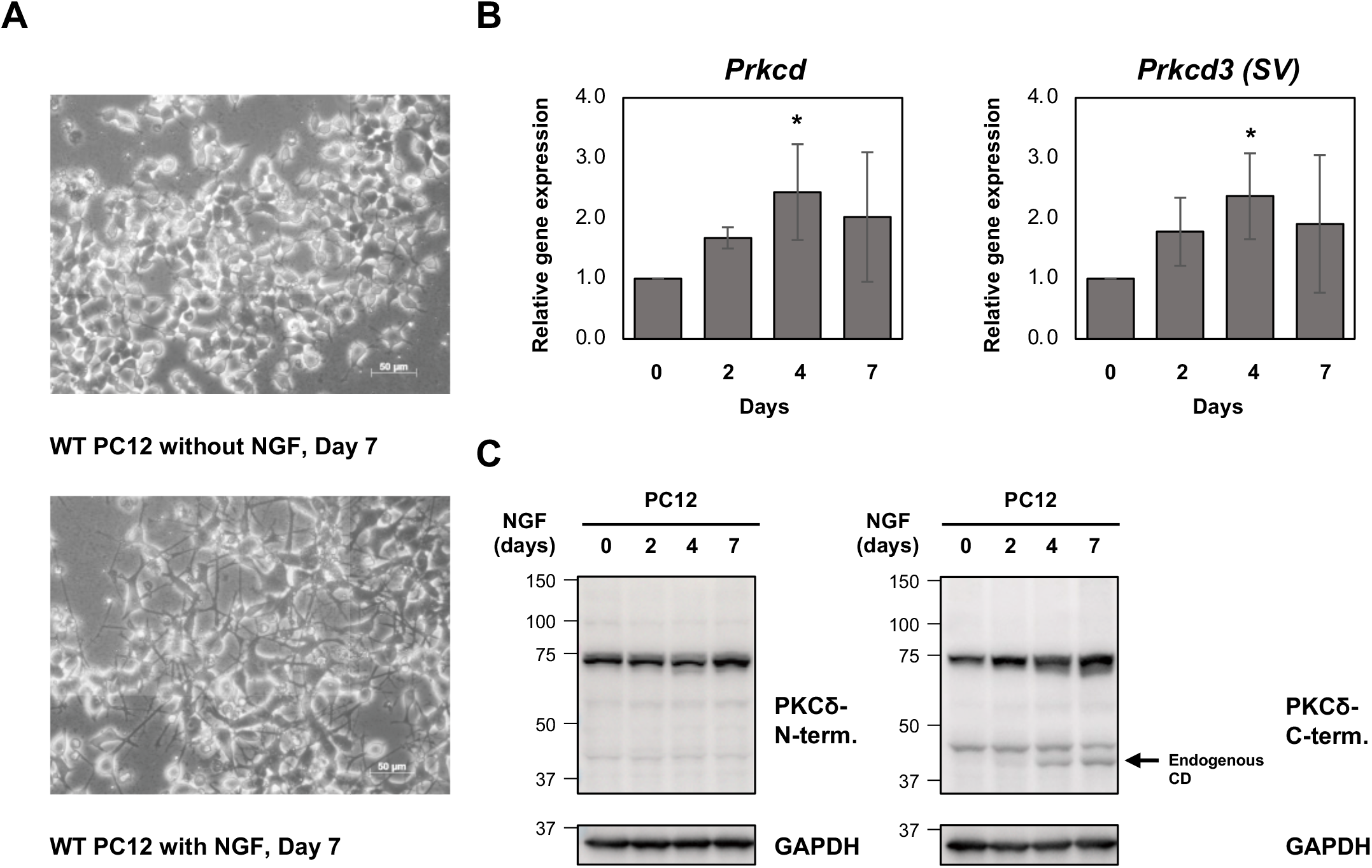
NGF treatment regulates PKCδ. (A) Representative images of neurite outgrowth induced by NGF treatment at day 7. (B) PC12 cells were cultured with or without 100ng/ml NGF. After 2, 4, and 7 days or before NGF treatment, total RNA were isolated. *Prkcd* and *Prkcd3 (SV)* transcripts were evaluated by qRT-PCR. Fold change was calculated upon normalization against *Gapdh* transcripts. *n* = 3. * *p*<0.05 (one-way ANOVA [Bonferroni’s test]). (C) PC12 cells were cultured with or without 100ng/ml NGF. After 2, 4, and 7 days, whole cell lysates were harvested. The cell lysates were subjected to immunoblotting analysis using the indicated antibodies. *n* = 2.

The protein levels of truncated PKCδ proteins were lower than that of PKCδ FL (Fig. 1B). This might be due to the instability of truncated PKCδ proteins. To determine whether truncated PKCδ RD, SV and CD are susceptible to their proteasomal degradation, we examined the effect of MG-132, which impairs proteasome activity, on the stability of truncated PKCδ RD, SV and CD. Ubiquitinated proteins definitely accumulated by the proteasome inhibition (Fig. 4A). Whereas PKCδ FL, RD, and CD did not accumulate by the proteasome inhibition, PKCδ SV did in a dose-dependent manner (Fig. 4B). These results suggested that truncated PKCδ SV was more susceptible to its proteasomal degradation than other PKCδ proteins were.

**Figure 4.**
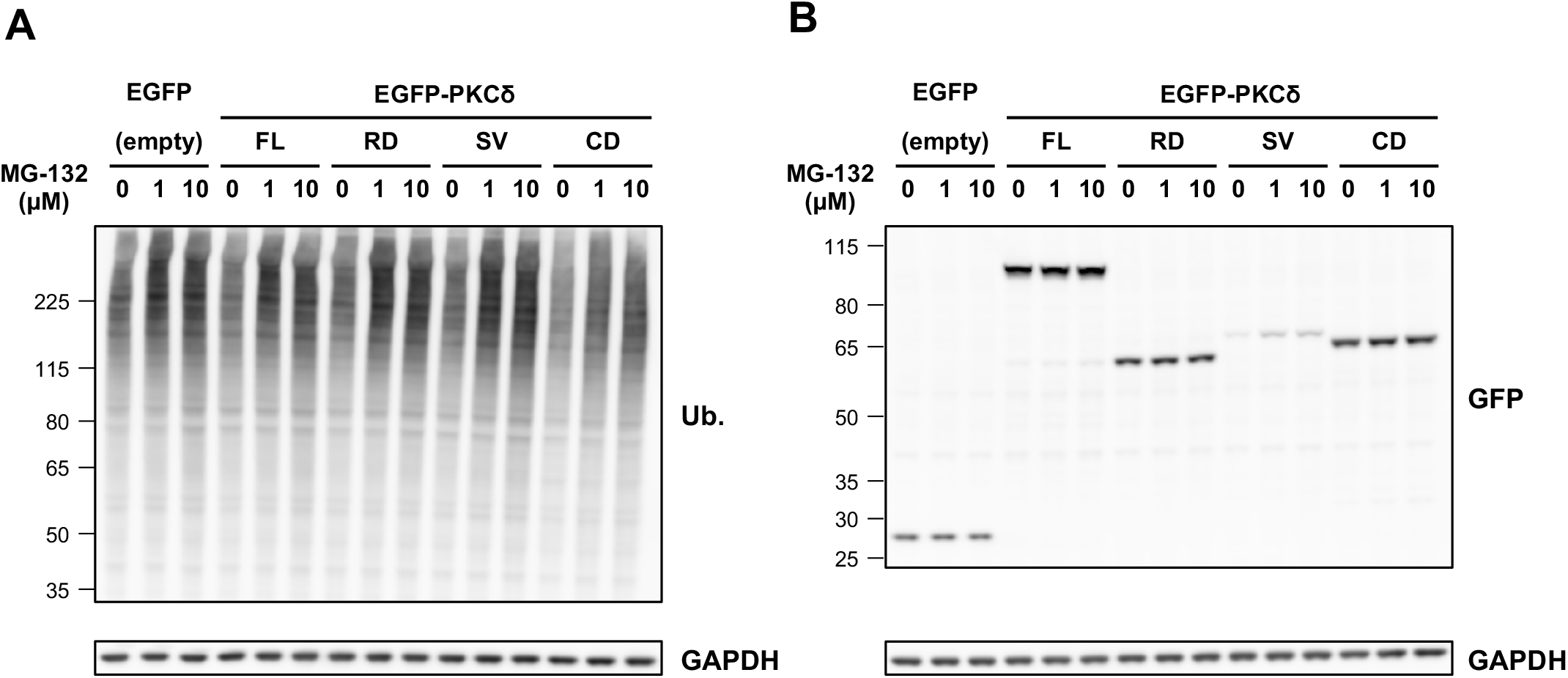
PKCδ SV is susceptible to its proteasomal degradation. (A and B) PC12 cells containing PKCδ constructs were cultured with 1ug/ml doxycycline. 22h later, the indicated concentration of MG-132 were added and cells were incubated for another 2h, and whole cell lysates were harvested. The cell lysates were subjected to immunoblotting analysis using the antibodies against GFP (A), ubiquitin (B), and GAPDH (loading control). *n* = 2.

## Discussion

PKCδ SV, also known as PKCδ III, was identified in rats and has been reported to localize in the cytoplasm and plasma membrane, but not in the nucleoplasm [12]. However, the physiological role of PKCδ SV remains unclear. In this study, we revealed a relationship between PKCδ SV and neurite outgrowth. The expression of PKCδ SV induced neurite outgrowth in PC12 cells without affecting endogenous ERK1/2 and PKCδ in the absence of NGF (Fig. 1 and 2). Conversely, PKCδ SV increased during NGF-mediated neurite outgrowth in mRNA level as well as PKCδ FL did (Fig. 3). These data emphasized the importance of PKCδ SV in neurite outgrowth.

During NGF-mediated neurite outgrowth, immunoblot analysis implied that the region recognized by anti-PKCδ (N term) was modified (Fig. 3C). The region recognized by anti-PKCδ (N term) contains two tyrosine residues (Y155 and Y187) in C1 domain that can be phosphorylated by tyrosine kinases. As PKC δ phosphorylation gradually decreased during NGF treatment (Fig. 2D), catalytic fragments appeared (Fig. 3C). In line with this observation, phosphorylated PKCδ is subject to be proteolytically degraded [18,19].

Although NGF-mediated neurite outgrowth depends on ERK activation [14,15], PKCδ RD- and SV-mediated neurite outgrowth was ERK-independent (Fig. 2). This implies the presence of the unique mechanisms by which PKCδ RD and SV induce neurite outgrowth. Neurite outgrowth is executed by actin and microtubule dynamics [1,20]. Both polymerization and depolymerization of actin and microtubule are important for neurite initiation [20]. Actin and microtubule dynamics might be regulated by PKCδ RD and SV. First, PKCδ was shown to inhibit platelet aggregation through its interaction with actin organizer VASP. PKCδ negatively regulates VASP through inhibiting its phosphorylation [21]. Interestingly, VASP is required for filopodia formation and subsequent neurite initiation during cortical development [22,23]. Furthermore, PKCδ RD functions as a dominant-negative mutant and inhibits noncanonical Wnt pathway and gastrulation movements in frogs [24]. Taken together, it is plausible that PKCδ RD and SV might prevent endogenous PKCδ FL from interacting with VASP, resulting in actin polymerization and neurite initiation in PC12 cells. Secondly, PKCδ interacts with Dishevelled (Dvl), and the activation of PKCδ is required for Dvl translocation to membrane [24]. Dvl mediates Wnt signaling to reorganize microtubule cytoskeleton by inducing APC loss from microtubule plus-ends [25]. PKCδ RD and SV might also prevent endogenous PKCδ FL from interacting with Dvl, resulting in APC binding to microtubule plus-ends and the inhibition of growth cone enlargement. Whilst current speculative, PKCδ RD and SV might regulate actin and microtubule polymerization. We also found that PKCδ SV was more susceptible to its proteasomal degradation than other PKCδ proteins were (Fig. 4). This observation raised the possibility that PKCδ SV would be utilized to fine tune the balance of polymerization and depolymerization.

Finally, we revealed that PKCδ SV induced neurite outgrowth in the absence of NGF in PC12 cells. PKCδ FL and SV increased their mRNA expression after NGF treatment. Our results provided the evidence that PKCδ SV potentially regulate neurite outgrowth during neuronal development. However, further studies are required to determine whether PKCδ SV is expressed in a neuronal developing brain and to elucidate the mechanisms by which PKCδ RD and SV induce neurite outgrowth.

## Acknowledgments

The authors are grateful to the staff of Support Center for Medical Research and Education, Tokai University for their excellent assistance with microscopies and their instructions for statistical data analysis and immunoblotting. This work was supported by 2020 Tokai University School of Medicine Research Aid.

## References

[1] J.S. Da Silva, C.G. Dotti, Breaking the neuronal sphere: Regulation of the actin cytoskeleton in neuritogenesis, Nat Rev Neurosci. 3 (2002) 694—704. 10.1038/nrn918.

[2] A.P. Barnes, F. Polleux, Establishment of axon-dendrite polarity in developing neurons, Annu Rev Neurosci. 32 (2009) 347—381. 10.1146/annurev.neuro.31.060407.125536.

[3] T. Takano, Y. Funahashi, K. Kaibuchi, Neuronal polarity: Positive and negative feedback signals, Front Cell Dev Biol. 7 (2019). 10.3389/fcell.2019.00069.

[4] S. Prem, J.H. Millonig, E. DiCicco-Bloom, Dysregulation of Neurite Outgrowth and Cell Migration in Autism and Other Neurodevelopmental Disorders, in: Adv Neurobiol, Springer, 2020: pp. 109–153. 10.1007/978-3-030-45493-7_5.

[5] J.D. Black, T. Affandi, A.R. Black, M.E. Reyland, PKCα and PKCδ: Friends and Rivals, Journal of Biological Chemistry. 298 (2022). 10.1016/j.jbc.2022.102194.

[6] M. Leitges, M. Mayr, U. Braun, U. Mayr, C. Li, G. Pfister, N. Ghaffari-Tabrizi, G. Baier, Y. Hu, Q. Xu, Exacerbated vein graft arteriosclerosis in protein kinase C*δ*-null mice, Journal of Clinical Investigation. 108 (2001) 1505—1512. 10.1172/JCI200112902.

[7] A. Miyamoto, K. Nakayama, H. Imaki, S. Hirose, Y. Jiang, M. Abe, T. Tsukiyama, H. Nagahama, S. Ohno, S. Hatakeyama, K.I. Nakayama, Increased proliferation of B cells and auto-immunity in mice lacking protein kinase C*δ*, Nature. 416 (2002) 865—869. 10.1038/416865a.

[8] I. Mecklenbräuker, K. Saijo, N.-Y. Zheng, M. Leitges, A. Tarakhovsky, Protein kinase Cδ controls self-antigen-induced B-cell tolerance, Nature. 416 (2002) 860—865. 10.1038/416860a.

[9] O. Bezy, T.T. Tran, J. Pihlajamäki, R. Suzuki, B. Emanuelli, J. Winnay, M.A. Mori, J. Haas, S.B. Biddinger, M. Leitges, A.B. Goldfine, M.E. Patti, G.L. King, C.R. Kahn, PKCδ regulates hepatic insulin sensitivity and hepatosteatosis in mice and humans, Journal of Clinical Investigation. 121 (2011) 2504–2517. 10.1172/JCI46045.

[10] W. Ma, C. Baumann, M.M. Viveiros, Lack of protein kinase C-delta (PKCδ) disrupts fertilization and embryonic development, Mol Reprod Dev. 82 (2015) 797–808. 10.1002/mrd.22528.

[11] Y.S. Niino, I. Kawashima, Y. Iguchi, H. Kanda, K. Ogura, K. Mita-Yoshida, T. Ono, M. Yamazaki, K. Sakimura, S. Yogosawa, K. Yoshida, S. Shioda, T. Gotoh, PKCδ deficiency inhibits fetal development and is associated with heart elastic fiber hyperplasia and lung inflammation in adult PKCδ knockout mice, PLoS One. 16 (2021). 10.1371/journal.pone.0253912.

[12] T. Ueyama, Y. Ren, S. Ohmori, K. Sakai, N. Tamaki, N. Saito, cDNA cloning of an alternative splicing variant of protein kinase C *δ* (PKC *δ*III), a new truncated form of PKCδ, in rats, Biochem Biophys Res Commun. 269 (2000) 557–563. 10.1006/bbrc.2000.2331.

[13] M. Ling, U. Trollér, R. Zeidman, C. Lundberg, C. Larsson, Induction of neurites by the regulatory domains of PKCδ and ε is counteracted by PKC catalytic activity and by the RhoA pathway, Exp Cell Res. 292 (2004) 135–150. 10.1016/j.yexcr.2003.08.013.

[14] D. Vaudry, P.J.S. Stork, P. Lazarovici, L.E. Eiden, Signaling Pathways for PC12 Cell Differentiation: Making the Right Connections, Science (1979). 296 (2002) 1648–1649. 10.1126/science.1071552.

[15] S. Cowley, H. Paterson, P. Kemp, C. Marshall, Activation of MAP kinase kinase is necessary and sufficient for PC12 differentiation and for transformation of NIH 3T3 cells, Cell. 77 (1994) 841–852. 10.1016/0092-8674(94)90133-3.

[16] K.R. O’Driscoll, K.K. Teng, D. Fabbro, L.A. Greene, I.B. Weinstein, Selective translocation of protein kinase C-delta in PC12 cells during nerve growth factor-induced neuritogenesis., Mol Biol Cell. 6 (1995) 449–458. 10.1091/mbc.6.4.449.

[17] K.C. Corbit, D.A. Foster, M.R. Rosner, Protein Kinase C*δ* Mediates Neurogenic but Not Mitogenic Activation of Mitogen-Activated Protein Kinase in Neuronal Cells, Mol Cell Biol. 19 (1999) 4209–4218. 10.1128/MCB.19.6.4209.

[18] J. Srivastava, K.J. Procyk, X. Iturrioz, P.J. Parker, Phosphorylation is required for PMA- and cell-cycle-induced degradation of protein kinase Cδ, Biochemical Journal. 368 (2002) 349–355. 10.1042/BJ20020737.

[19] Z. Lu, T. Hunter, Degradation of activated protein kinases by ubiquitination, Annu Rev Biochem. 78 (2009) 435–475. 10.1146/annurev.biochem.013008.092711.

[20] R. Sainath, G. Gallo, Cytoskeletal and signaling mechanisms of neurite formation, Cell Tissue Res. 359 (2014) 267–278. 10.1007/s00441-014-1955-0.

[21] G. Pula, K. Schuh, K. Nakayama, K.I. Nakayama, U. Walter, A.W. Poole, PKCδ regulates collagen-induced platelet aggregation through inhibition of VASP-mediated filopodia formation, Blood. 108 (2006) 4035–4044. 10.1182/blood-2006-05-023739.

[22] E.W. Dent, A. V. Kwiatkowski, L.M. Mebane, U. Philippar, M. Barzik, D.A. Rubinson, S. Gupton, J.E. Van Veen, C. Furman, J. Zhang, A.S. Alberts, S. Mori, F.B. Gertler, Filopodia are required for cortical neurite initiation, Nat Cell Biol. 9 (2007) 1347–1359. 10.1038/ncb1654.

[23] A. V. Kwiatkowski, D.A. Rubinson, E.W. Dent, J. Edward van Veen, J.D. Leslie, J. Zhang, L.M. Mebane, U. Philippar, E.M. Pinheiro, A.A. Burds, R.T. Bronson, S. Mori, R. Fässler, F.B. Gertler, Ena/VASP Is Required for Neuritogenesis in the Developing Cortex, Neuron. 56 (2007) 441–455. 10.1016/j.neuron.2007.09.008.

[24] N. Kinoshita, H. Iioka, A. Miyakoshi, N. Ueno, PKCδ is essential for Dishevelled function in a noncanonical Wnt pathway that regulates Xenopus convergent extension movements, Genes Dev. 17 (2003) 1663–1676. 10.1101/gad.1101303.

[25] S.A. Purro, L. Ciani, M. Hoyos-Flight, E. Stamatakou, E. Siomou, P.C. Salinas, Wnt regulates axon behavior through changes in microtubule growth directionality: A new role for adenomatous polyposis coli, Journal of Neuroscience. 28 (2008) 8644–8654. 10.1523/JNEUROSCI.2320-08.2008.

